# Isolation and characterisation of Lytic Jumbo Klebsiella phage BMPCR_005026

**DOI:** 10.64898/2026.09.30.755614

**Authors:** Isla Lloyd-Evans, Matthew Ford, Jed Hazzard, Casey Bartis, Faizal Patel, Navodya Roemer, Kayne Wright, Kane Porter, Max Coleman, Edouard Galyov, Spyridon Megremis, Martha R.J. Clokie, Andrew D. Millard, Slawomir Michniewski, Rachael C. Wilkinson

**Author notes:** **Author Disclosure statement:** No competing financial interests exist for any authors.

## Abstract

**Background:** *Klebsiella pneumoniae* is a key causative agent of multidrug resistant nosocomial infections. Bacteriophages offer a promising alternative treatment, but the diversity of *Klebsiella* species requires the characterisation of a broad range of phages, particularly the poorly characterised jumbo phages.

**Materials and Methods:** Phage BMCPR_005026 was isolated from sewage against *Klebsiella pneumoniae*. Whole genome sequencing, transmission electron microscopy and host range analysis against a diverse set of 40 clinically relevant *Klebsiella* strains were used to characterise the phage.

**Results:** The virion had a myovirus morphology and genomic analysis placed it within an undefined family in *Caudoviricetes* where it forms a new species within the genus *Alcyoneusvirus*. BMCPR_005026 could form plaques on 3 of the 40 *Klebsiella* strains tested (7.5%) and formed halos.

**Conclusion:** BMCPR_005026 represents a new species of phage within the genus *Alcyoneusvirus* with activity against 3 *Klebsiella* KL-types.

## Introduction

*Klebsiella pneumoniae* is a gram-negative, non-motile bacterium and a major cause of nosocomial infections, including urinary tract infections (UTIs), pneumonia, septicaemia, meningitis, and soft tissue infections^1–5^. *K. pneumoniae* is classed as a critical-priority antibiotic-resistant bacterium by WHO, surpassing methicillin-resistant *Staphylococcus aureus* (MRSA) in factors such as treatability, epidemiological global burden, and increase in drug resistance^6, 7^. Antimicrobial resistance (AMR) is an ever-increasing global threat, with an estimated 8.22 million deaths associated with AMR predicted to occur annually by 2050^8^. The development of new antimicrobial strategies will therefore be important in reducing the future burden of AMR-associated deaths^9^

The limited development of new small-molecule antibiotics has driven an increased interest in the potential of bacteriophages (phages), viruses which infect bacteria, as an alternative approach in the treatment of classical and hypervirulent *K. pneumoniae* infections^4,10^. One of the key advantages of phages is their host specificity; consequently, the isolation and characterisation of genetically diverse phages is essential for the development of effective phage cocktails. To date, there have been 2387 *Klebsiella* phages isolated and annotated, with only 31 classed as jumbo phages (genome > 200 kb)^11, 12^.

Jumbo phages are distinguished not only in their size, but also in their organisation at the genomic level and virion structure, while their evolutionary origin remains largely unknown^13, 14^. Jumbo phages can contain unique features that facilitate infection of their bacterial host, with some encoding their own molecular machinery and others producing a pseudo-nucleus shell to protect their genomes^14–16^. Furthermore, multiple *Klebsiella* jumbo phages have demonstrated a broad host range due to their expanded genomes and reduced dependence on the host proteins^13^. Characterisation of new jumbo phages is therefore important for expanding our understanding of phage diversity, genome organisation and mechanisms of host infection, whilst potentially identifying features relevant to the development of phage therapeutics. Here we present the characterisation of a novel *Klebsiella pneumoniae* jumbo phage BMCPR_005026.

## Materials and methods

Unless stated otherwise, all plasticware was purchased from SARSTEDT (Germany). All strains were grown in Luria-Bertani (LB) Miller media (Sigma Aldrich, Germany). Agar plates (bottom agar) were made at 1.5% w/v and all top agar at 0.45% w/v. All filter steps used 0.45 µm pore-size syringe filters.

### Bacterial growth conditions

The propagation strain *Klebsiella pneumoniae* LEI0296 was obtained from a rectal swab from the Leicester Royal Infirmary. Bacterial host range strains were selected from the BMCPR phage biobank collection, University of Leicester. Bacterial cultures were freshly streaked onto LB plates and incubated overnight at 37 °C. Overnight cultures of host strain LEI0296 were set up from a single colony and grown at 37 °C, 180 rpm for 18 hours, before inoculating LB broth (1 in 100 dilution) and grown at 37 °C, 180 rpm for 2.5 hours.

### Isolation of phage BMCPR_005026

Plaque assays were performed as described previously^17^, with *K. pneumoniae* LEI0296 as the host, with filtered sewage from Barston, UK. A resultant single plaque was picked using a 200 µL filter tip into 100 µL of sterile SM buffer (100 mM NaCl, 8 mM MgSO_4_ •7H_2_O, 50 mM Tris-HCl, pH 7.5). The plaque pick was confirmed as phage by the observation of a clearing zone, as described previously via the direct spot assay method with 10 µL spots^18^. The pick was then confirmed to be a single phage using the Plaque-2-seq approach^19^. The phage lysate was stored at 4 °C prior to further use.

A working phage stock was produced by infecting mid-exponential LEI0296 culture (10 mL; OD600 nm ∼ 0.4) with 20 µL of phage for 4 hours at 37 °C with shaking. The lysate was centrifuged (3392 *×g*, 15 minutes) and filtered through a 0.45 µm syringe filter.

High titre stocks for imaging analysis were made using an accumulative plate wash method. Plaque assays (5 total) were set up with 100 µL of working stock phage and 100 µL of LEI0296 to produce confluent lysis. An accumulative plate wash was performed by adding 8 mL of SM buffer to one plaque assay plate, left rocking at room temperature for 10 minutes. The top agar and SM were scraped up using an L-shaped cell spreader (Fisher Scientific, UK) and the mixture was poured onto the next plate, then left rocking at room temperature for 10 minutes. The process was repeated across five plates. The final mixture was centrifuged at 3392 *×g* for 15 minutes and the supernatant filtered. The final phage titre was determined via direct spot assay as above.

### Host range assay of phage BMCPR_005026

40 clinical *Klebsiella* strains with varying capsule types and sequence types were selected from the BMCPR phage biobank, University of Leicester. Bacterial strains were grown overnight as above and 300 µL of each was added to 8 mL of LB top agar and poured onto square LB bottom agar plates (Greiner Bio-One Ltd., Stonehouse, UK) and left to solidify. A ten-fold serial dilution to 10^−10^ of BMPCR_005026 was made, and 5 µL of each dilution spotted in triplicate. Once spots were dry, plates were incubated overnight at 37 °C. Plaques were counted and efficiency of plating (EOP) was calculated as a percentage (average PFU on target bacteria/ average PFU on host bacteria) along with the standard deviations for each strain.

### Genome sequencing, assembly and annotation of phage BMCPR_005026

BMCPR_005026 genome was sequenced according to the Plaque-2-seq method^19^. Briefly, an aliquot of the sample underwent DNaseI (Thermo Scientific, UK) treatment to remove host DNA, heat treatment to inactivate the DNaseI and release the DNA from virions. DNA was then amplified using EquiPhi29™ DNA Amplification Kit (Thermo Scientific, UK) followed by debranching using S1 nuclease (Thermo Scientific, UK). DNA was sequenced via Oxford Nanopore Technologies (ONT) using the Rapid sequencing DNA V14 - barcoding kit (SQK-RBK114.96) on a GridION, with the R10.4.1 flow cell. Dorado v7.3.9 with model dna_r10 4.1_e8 2_400bps_hac v4 3.0, was used for basecalling. Long reads were converted to short reads of length 300 bp, normalized and assembled with SPAdes v3.13.1. Assembled genomes underwent CheckV^20^ analysis, taxonomy assigned using *taxMyPhage*^21^ and lifestyle and safety assessed with PhageLeads^22^.

Phage genomes were reordered based on the first intergenic region upstream of the terminase large subunit, with genes being predicted and annotated using Prokka^23^ with PHROGs^24^ database.

### Purification of phage BMCPR_005026 for TEM

BMCPR_005026 was purified using ammonium acetate. 5 mL high titre (9.5 × 10^9^ PFU mL^-1^) phage lysate was added to 25 mL SM buffer and centrifuged at 20,000 *×g* for 1 hour. The phage pellet was resuspended with 30 mL 0.1 M ammonium acetate (Sigma-Aldrich, Germany) and centrifuged again at 20,000 *×g* for 1 hour. The final pellet was resuspended in 200 µL SM buffer. Titre was determined using direct spot assays as detailed above.

### Transmission Electron Microscopy (TEM)

5 µL of purified phage BMCPR_005026 (3.5 × 10^10^ PFU mL^-1^) was applied to a freshly glow discharged carbon film copper grid. This grid was glow discharged in a GloQube Plus (Quorum Technologies) for 30 seconds at 30 mA. Once applied, the sample was left to adsorb for 2 minutes. The excess liquid was blotted off with filter paper, before washing the grid twice with ultrapure water and staining with 5 µL of 1% (w/v) uranyl acetate. The sample was viewed on a JEOL JEM-1400 TEM with a voltage of 120 kV. Digital images were collected with an EMSIS

Xarosa digital camera with Radius software. ImageJ software (Version 1.54g) was used to calculate lengths and averages were taken from measurements across ten phage particles^25^.

## Results

Phage BMCPR_005026 (accession number: OZ545911) was isolated from a sewage sample from Barston, West Midlands, UK against clinical *Klebsiella pneumoniae* strain isolate LEI0296.

TEM analysis of BMCPR_005026 demonstrated a myovirus morphology, with a total virion length of approximately 260 ± 8.6 nm (Figure 1a). Well-preserved structural features such as the icosahedral capsid head, measuring approximately 123 ± 3.0 nm (width) by 136 ± 3.9 nm (length), and contractile tail, measuring approximately 124 ± 4.2 nm (length) were observed. BMCPR_005026 produces small plaques (0.2 ± 0.04 mm) on the host strain with large halo clearances 1.8 ± 0.36 mm in diameter after 18 hours incubation at 37 °C (Figure 1b).

**Figure 1.**
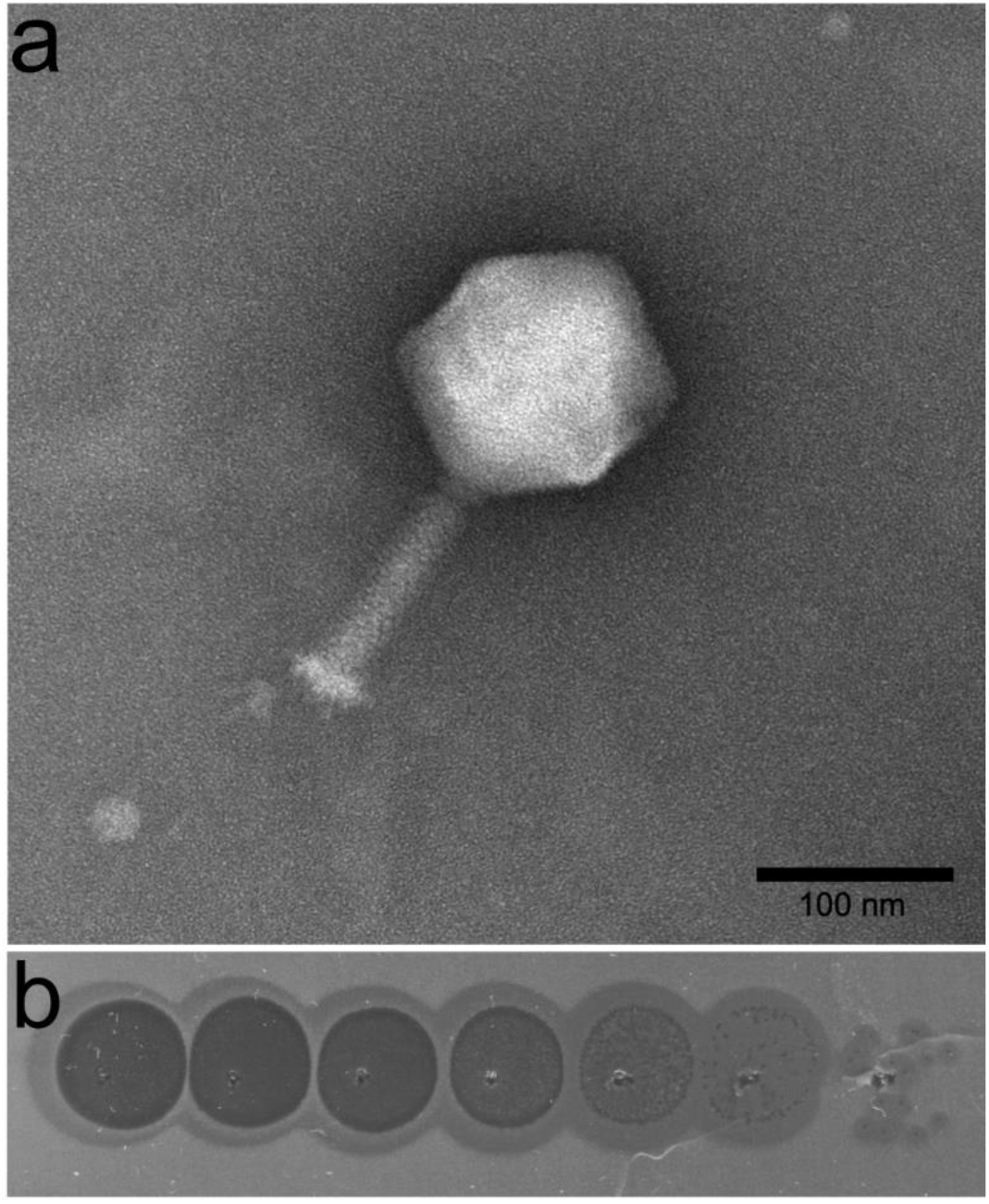
Characterisation of phage BMCPR_005026. a) Morphology of phage BMCPR_005026. Virions were imaged by TEM after negative staining with uranyl acetate, at 5000x magnification. b) Scanned image of direct spot assays for a 10-fold dilution series of purified BMCPR_005026 on original host LEI0296 from 10^−1^ to 10^−7^ PFU mL^-1^, with halos visible at all dilutions.

Genomic sequencing of BMCPR_005026 revealed a genome of 352,398 bp in length with a GC content of 32% and 544 predicted genes, of which four are tRNAs. Analysis with PhageLeads^22^ identified no known temperate markers, antimicrobial resistance (AMR) genes, or virulence genes. Only 31.25% of genes could be ascribed a predicted function using PHROGs (Supplementary Table S1).

Initial blastn analysis identified that BMCPR_005026 had significant similarity to five phages. Subsequent analysis of all five phages with *taxMyPhage* identified BMCPR_005026 as a new species within the genus *Alcyoneusvirus* (Figure 2), in addition to Klebsiella phage vB_KleM_RaK2 (accession: JQ513383)^27^ and Klebsiella phage K64-1 (accession: AB897757)^28^ which are already assigned taxonomy by ICTV^26^. The currently unclassified phages from blastn analysis, Muenster (Accession: MT708547) and KP13-7 (Accession: OP617745), also lie within this genus. All five phages are separate species within the genus *Alcyoneusvirus*, thus, expanding the number of species in the genus *Alcyoneusvirus* from two to five. Further phylogenetic analysis revealed members of the genus *Alcyoneusvirus* form a deep branching clade that is a sister group to the phages of the well-known ‘jumbo’ genus *Asteriusvirus* (Figure 3).

**Figure 2.**
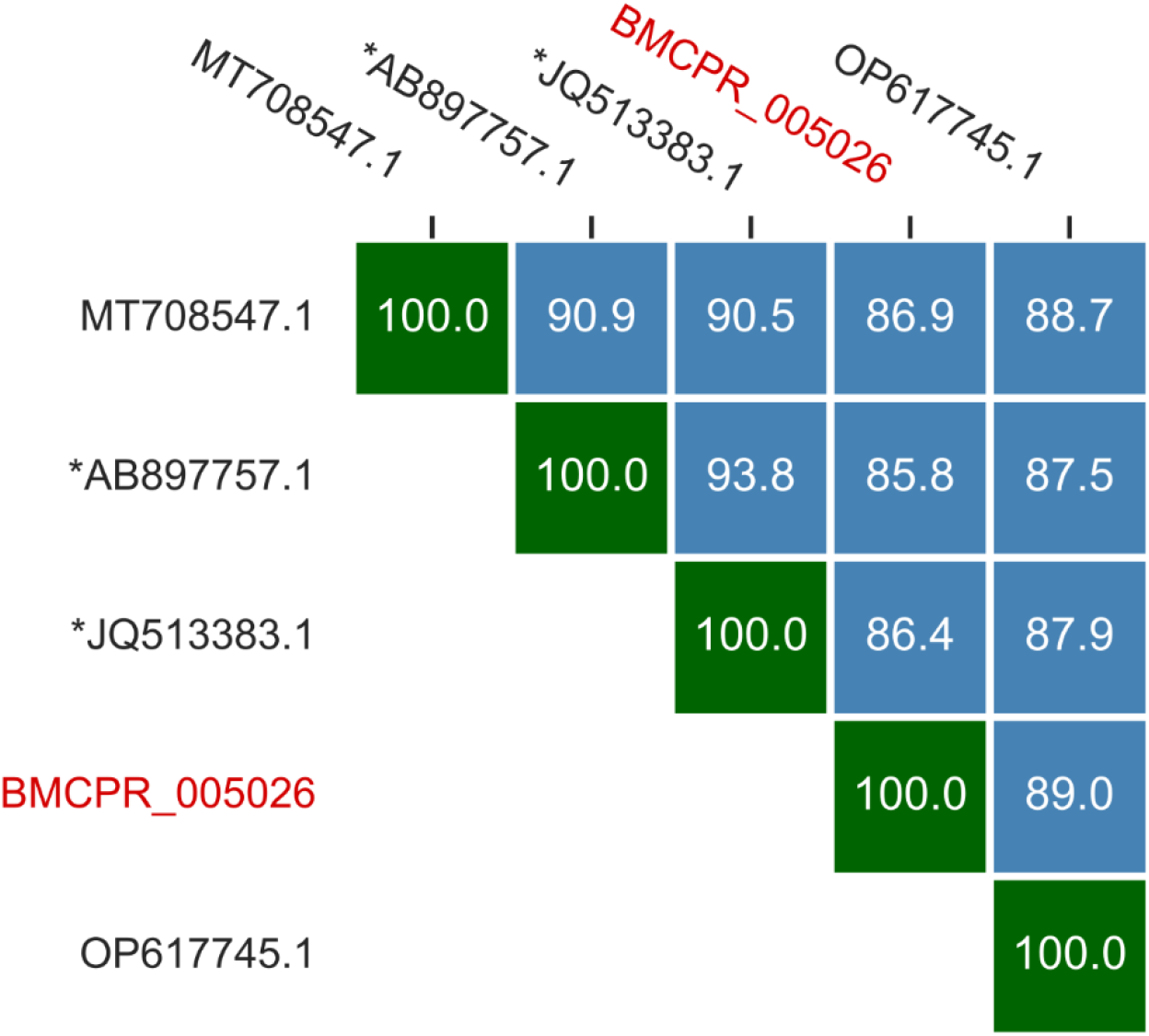
Classification of BMCPR_005026 (red) within the genus *Alcyoneusvirus*. Upper right similarity matrix generated by *taxMyPhage* of phage BMCPR_005026 along with MT708547.1 (Muenster); AB897757.1 (K64-1); JQ513383.1 (vB_KleM_RaK2); and OP617745.1 (KP13-7). BMCPR_005026 had <95% ANI with current phage species of the genus *Alcyoneusvirus* (denoted with *), *Alcyoneusvirus K64-1* (AB897757.1) and *Alcyoneusvirus RaK2 (*JQ513383.1). BMCPR_005026 had > 70% ANI with all phages.

**Figure 3.**
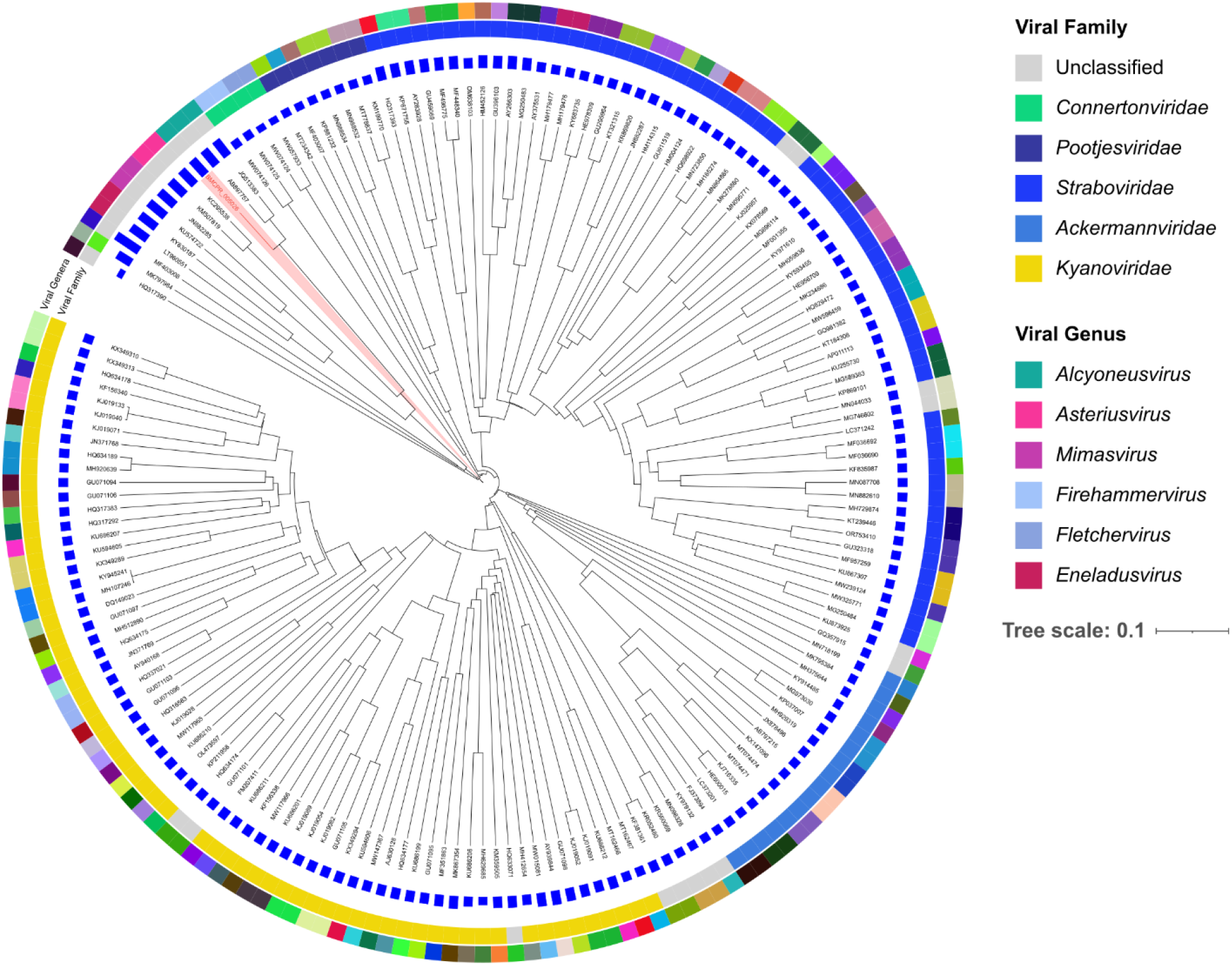
Proteomic phylogeny of phage BMCPR_005026. Phylogenetic placement of BMCPR_005026 (highlighted in red) was assessed using a proteome-based viral tree generated with ViPTreeGen (PMID: 28379287). Representatives from all defined genera within the *Caudoviricetes* along with related phages lacking formal taxonomy, obtained from INPHARED2, were included for analysis (n=2800). The tree was visualised and pruned in iTOLVersion 7.6 (PMID 38613393), to allow visualisation of the closest related families and genera. Current members of the *Alcyoneusvirus* genus vB_KleM_RaK2 (JQ513383) and K64-1 (AB897757) are included. The outer ring is the Viral Family and the inner ring Viral Genera. Genome size is indicated by the blue bars to the inner of the classification rings.

Host range analysis against 40 clinically relevant *Klebsiella spp*. strains representing 36 distinct KL-types, revealed only three were permissive, all of which were *K. pneumoniae* (7.5%) (Table 1). These comprised the original isolation host, LEI0296 (KL60), and two additional strains, LEI0372 (KL136) and LEI0476 (KL30). Large ‘halos’ as seen in Figure 1b, could be observed with BMCPR_005026 across all 3 KL-types. The efficiencies of plating (EOP) against LEI0372 and LEI0476 were 229.56% and 2.15%, respectively. Non-permissive strains and K-types can be found in Supplementary Table S2.

**Table 1:** Lytic activity of BMCPR_005026 against various *Klebsiella* strains. Only permissive strains are shown here. ^1^ST, sequence type. ^2^EOP = Efficiency of plating, percentage calculated: 100 x (PFU mL^-1^ on strain / PFU ml^-1^ on original host).

| Bacterial strain | Species | ST<sup>1</sup> | K locus | PFU m<sup>-1</sup> | EOP<sup>2</sup> |
| --- | --- | --- | --- | --- | --- |
| *LEI0296 (host)* | *Klebsiella pneumoniae* | 1658 | KL60 | 1.80E+05 | 100.00% |
| *LEI0372* | *Klebsiella pneumoniae* | 1419 | KL136 | 4.13E+05 | 229.56% |
| *LEI0476* | *Klebsiella pneumoniae* | 234 | KL30 | 3.87E+03 | 2.15% |

## Discussion

With the rise in multidrug resistant *Klebsiella* infections, the isolation of new phages active against clinically relevant strains is vital. Here we have described the isolation and characterisation of a new jumbo phage within the genus *Alcyoneusvirus*, phage BMCPR_005026, with activity against three *Klebsiella* KL-types.

Through comparative genomic analysis we have identified several species within the genus *Alcyoneusvirus*, within which we have identified phage BMCPR_005026 as being representative of a new viral species. We propose the species name “Alcyoneusvirus barston*”*. Genomic analysis demonstrated a genome of 352.4 kb with 544 predicted ORFs, 68% of which were annotated as hypothetical. Although this proportion of hypothetical proteins is considerably high, it represents a smaller percentage than that observed in some of the other *Klebsiella* phages. For comparison, phage vB_KpnM_FBKp24 contains 79% hypothetical proteins, while phage Miami contains 84.3%^29,30^. Genes encoding putative additional metabolic functions, often associated with other lytic jumbo phages, include the presence of genes associated with NAD+ salvage, heavy metal (specifically tellurite resistance) and proteolytic complexes^31–33^. The genes associated with tellurite resistance have previously been found in coliphages of the genus *Asteriusvirus*, which was found to be the closest related genera of phages to the *Alcyoneusviruses*. Suggesting that these genes may represent an evolutionarily conserved feature retained since the divergence of these related phage lineages. If resistance to tellurite is conferred or they serve some other function, remains to be determined.

The phage ΦK64-1, another jumbo phage from the *Alcyoneusvirus* genus, has been shown to infect 11 different K types^34^. Here we found that BMCPR_005026 displayed a narrow host range determined by KL-type, infecting only 3 of 40 *Klebsiella spp*. strains tested (7.5%), all three of which were *K. pneumoniae*. It is worth noting that of the 11 K-types permissive to ΦK64-1, BMCPR_005026 was only able to infect KL30, suggesting large variation within the phage genus, however we were unable to test against 3 of the 11 K-types^34^. BMCPR_005026 is therefore consistent with what has been observed more broadly for other *K. pneumoniae* phages, demonstrating infectivity between 1% and 31% of hosts tested, which is thought to link in part to the diverse nature of capsule types seen across *Klebsiella spp*.^35, 36^. This is further highlighted by Ferriol-González *et al*. who screened for the cross-reactivity of 86 isolated *K. pneumoniae* phages against 77 *K. pneumoniae* strains and demonstrated that each phage on average only infected 2.67 strains, which is consistent with the three found for BMCPR_005026^37^. Interestingly, the isolation strain *K. pneumoniae* LEI0296 has the KL60 type, which correlates with the K60 capsule. This capsule structure is distinct as it features three glucopyranosyl side chains per heptasaccharide subunit, in contrast to the one or two side chains typical of most other *Klebsiella* capsules^38^. The bulkier nature of this capsule type makes the susceptibility of this strain of particular interest. Capsule complexity, however, did not correlate with genomic profiles of other permissible *K. pneumoniae* strains. For instance, strain LEI0476 has the KL30 type, which correlates to a K30 capsule known to have two side chains per unit^39^. Further comparison with strain LEI0392, with the KL136 type, was not possible as the capsule structure, to the best of our knowledge, has yet to be determined.

In summary, the genomic characterisation and classification of phage BMCPR_005026 as a member of the proposed species “*Alcyoneusvirus barston”* expands the available genomic resource for *Klebsiella* jumbo phages. These data provide a valuable reference for future studies exploring jumbo phage genomic diversity and the complex mechanisms driving capsule-mediated host susceptibility.

## Acknowledgements

Bioinformatics analysis was carried out using MRC CLIMB Infrastructure MR/L015080/1. RW, MF, SMich, FP, KP, KW, ILE and NR were funded by philanthropic support to create the Becky Mayer Centre for Phage Research. MCol was funded by the College of Life Science, University of Leicester. Sample processing and TEM analysis was completed within the Electron Microscopy Facility, University of Leicester. All sample processing and imaging was done by Natalie Allcock with all consumables acquired through Agar Scientific.

