## Supplemental Tables for "Isolation and characterisation of Lytic Jumbo Klebsiella phage BMPCR_005026"

**Supplementary Table S1.** Table displaying the number of genes in each category according to the PHROGs database (PMID 34377978).

| **Category** | **Number of genes** |
| --- | --- |
| Head and packaging | 42 |
| DNA, RNA and nucleotide metabolism | 60 |
| Moron, auxiliary metabolic gene, host takeover | 10 |
| Transcription regulation | 3 |
| Tail | 24 |
| Lysis | 3 |
| Head-to-Tail Connectors | 2 |
| Other | 26 |
| Unknown function (hypothetical) | 374 |
| tRNA genes | 4 |
| **Total** | **548** |

**Supplementary Table S2.** Lytic activity of BMCPR_005026 against various *Klebsiella* strains of multiple species. ^1^ST, sequence type. ^2^Lysis: (1) - lytic activity observed, (0) - no lytic activity observed. ^3^EOP = Efficiency of plating, percentage calculated: 100 x (PFU ml^-1^ on strain / PFU ml^-1^ on original host).

| **No.** | **Bacterial strain** | **Species** | **ST^1^** | **K locus** | **Lysis^2^** | **PFU ml^-1^** | **EOP^3^** |
| --- | --- | --- | --- | --- | --- | --- | --- |
| 1 | **LEI0296 (host)** | ***Klebsiella pneumoniae*** | **1658** | **KL60** | **1** | **1.80E+05** | **100.00%** |
| 2 | LEI0024 | *Klebsiella pneumoniae* | 180 | KL54 | 0 |  |  |
| 3 | LEI0028 | *Klebsiella variicola subsp. variicola* | 906 | KL10 | 0 |  |  |
| 4 | LEI0240 | *Klebsiella pneumoniae* | 147 | KL10 | 0 |  |  |
| 5 | LEI0254 | *Klebsiella pneumoniae* | 147 | KL48 | 0 |  |  |
| 6 | LEI0278 | *Klebsiella pneumoniae* | 1537 | KL124 | 0 |  |  |
| 7 | LEI0291 | *Klebsiella pneumoniae* | 29 | KL19 | 0 |  |  |
| 8 | LEI0361 | *Klebsiella pneumoniae* | 466 | KL22 | 0 |  |  |
| 9 | LEI0363 | *Klebsiella quasipneumoniae subsp. similipneumoniae* | 414 | KL123 | 0 |  |  |
| 10 | LEI0364 | *Klebsiella pneumoniae* | 133 | KL47 | 0 |  |  |
| 11 | LEI0365 | *Klebsiella pneumoniae* | 133 | KL47 | 0 |  |  |
| 12 | LEI0366 | *Klebsiella pneumoniae* | 45 | KL62 | 0 |  |  |
| 13 | **LEI0372** | ***Klebsiella pneumoniae*** | **1419** | **KL136** | **1** | **4.13E+05** | **229.56%** |
| 14 | LEI0390 | *Klebsiella pneumoniae* | 1138 | KL158 | 0 |  |  |
| 15 | LEI0392 | *Klebsiella pneumoniae* | 1613 | KL12 | 0 |  |  |
| 16 | LEI0397 | *Klebsiella pneumoniae* | 2188 | KL136 | 0 |  |  |
| 17 | LEI0403 | *Klebsiella pneumoniae* | 6260 | KL17 | 0 |  |  |
| 18 | LEI0404 | *Klebsiella pneumoniae* | 147 | KL64 | 0 |  |  |
| 19 | LEI0409 | *Klebsiella pneumoniae* | 116 | KL11 | 0 |  |  |
| 20 | LEI0419 | *Klebsiella pneumoniae* | 307 | KL02 | 0 |  |  |
| 21 | LEI0428 | *Klebsiella variicola subsp. variicola* | 347 | KL9 | 0 |  |  |
| 22 | LEI0431 | *Klebsiella pneumoniae* | 3072 | KL39 | 0 |  |  |
| 23 | LEI0437 | *Klebsiella pneumoniae* | 39 | KL2 | 0 |  |  |
| 24 | LEI0440 | *Klebsiella pneumoniae* | 17 | KL23 | 0 |  |  |
| 25 | LEI0445 | *Klebsiella pneumoniae* | 7498 | KL113 | 0 |  |  |
| 26 | LEI0451 | *Klebsiella quasipneumoniae subsp. similipneumoniae* | 334 | KL103 | 0 |  |  |
| 27 | LEI0453 | *Klebsiella pneumoniae* | 70 | KL122 | 0 |  |  |
| 28 | LEI0456 | *Klebsiella pneumoniae* | 2703 | KL16 | 0 |  |  |
| 29 | LEI0460 | *Klebsiella pneumoniae* | 2494 | KL21 | 0 |  |  |
| 30 | LEI0468 | *Klebsiella pneumoniae* | 1726 | KL81 | 0 |  |  |
| 31 | LEI0469 | *Klebsiella pneumoniae* | 726 | KL131 | 0 |  |  |
| 32 | LEI0473 | *Klebsiella pneumoniae* | 45 | KL5 | 0 |  |  |
| 33 | **LEI0476** | ***Klebsiella pneumoniae*** | **234** | **KL30** | **1** | **3.87E+03** | **2.15%** |
| 34 | LEI0479 | *Klebsiella pneumoniae* | 1507 | KL114 | 0 |  |  |
| 35 | LEI0481 | *Klebsiella pneumoniae* | 1411 | KL61 | 0 |  |  |
| 36 | LEI0491 | *Klebsiella pneumoniae* | 45 | KL24 | 0 |  |  |
| 37 | LEI0494 | *Klebsiella pneumoniae* | 502 | KL15 | 0 |  |  |
| 38 | LEI0500 | *Klebsiella pneumoniae* | 1401 | KL49 | 0 |  |  |
| 39 | LEI0509 | *Klebsiella pneumoniae* | 133 | KL116 | 0 |  |  |
| 40 | LEI0521 | *Klebsiella pneumoniae* | 1552 | KL25 | 0 |  |  |
